# Proximity proteomics reveals a role for IFI16 during human coronavirus infection

**DOI:** 10.64898/2026.04.22.720112

**Authors:** Sylvester Languon, Ian Bailey, Madeleine Sorensen, Zachary D. Miller, Jihui Sha, Jake Dearborn, William Dowell, Tylar Kirch, James Wohlschlegel, Devdoot Majumdar

## Abstract

Viruses rely on the infected host cell to ensure successful replication and propagation of infection. This is achieved through interactions between virus-encoded proteins and proteins expressed in infected cells. All human coronaviruses (HCoVs) encode 16 non-structural proteins (NSPs) which exhibit some level of similarity in identity and function among the HCoVs. To identify host proteins that are potential interacting partners of HCoV NSPs, we utilized split-TurboID along with mass spectrometry and identified IFN-γ-inducible protein-16 (IFI16) as a proximal partner of SARS-CoV-2 NSP8 and NSP10. To investigate the significance of the association between the NSP8/NSP10 complex and IFI16, we utilized CRISPR-Cas9 to knockout and CRISPRi to knockdown IFI16 in A549 cells and demonstrated that loss or reduced expression of IFI16 leads to a decrease in human coronavirus infection. We further demonstrated that there is reduced viral RNA replication and viral protein synthesis upon loss of IFI16. Interestingly, the loss of IFI16 results in reduced expression of type I IFNs. Taken together, these data suggests that IFI16 promotes human coronavirus infection, and the role IFI16 plays in coronavirus replication is independent of its role as a regulator of type I IFN gene expression.

## Introduction

Coronaviruses are enveloped, positive-sense single-stranded RNA viruses that are classified under the family *Coronaviridae* which consist of four genera *Alpha-, Beta-, Gamma-*, and *Deltacoronavirus*. While Alpha- and betacoronaviruses are known to infect several mammalian species, gamma- and deltacoronaviruses have been found to infect avian species(1,2). Coronaviruses possess a non-segmented genome of ∼30-kb and this encodes 16 non-structural proteins (nsp1–nsp16), four or five structural proteins [spike (S), membrane (M), envelope (E), and nucleocapsid (N), hemagglutinin-esterase (HE)], and a variety of accessory proteins depending on the subgenera of the virus (3,4). Currently, two alphacoronaviruses [Human coronavirus (HCoV)-229E and HCoV-NL63] and five betacoronaviruses [HCoV-OC43, HCoV-HKU1, severe acute respiratory syndrome (SARS)-CoV, Middle East respiratory syndrome (MERS)-CoV and SARS-CoV-2] have been found to infect humans. Of the seven coronaviruses that infect humans, HCoV-OC43, HCoV-HKU1, (HCoV)-229E and HCoV-NL63 cause common cold symptoms and are considered endemic coronaviruses due to their continuous circulation in the human population(5,6). In contrast, MERS-CoV, SARS-CoV and SARS-CoV-2 are highly pathogenic and have caused severe respiratory disease outbreaks over the last 20 years, due to spillover events from other mammalian host species into the human population(7,8). The occurrence of multiple coronavirus outbreaks over the past 2 decades underscores the need for research to understand coronavirus biology and the virus-host interactions that occur during infection, with the goal of identifying targets for the design of drugs and production of vaccines.

Several proteomic approaches have been employed to identify the interacting partners of SARS-CoV-2 encoded proteins, with the goal of identifying protein targets for the design and evaluation of host-directed therapies against SARS-CoV-2. These approaches include affinity purification mass spectrometry (AP-MS) (9–11,11–13), cross-linking MS (XL-MS)(14,15), and proximity dependent labelling MS (PDL-MS) (16–18). The utilization of these methods generated a wealth of information about how SARS-CoV-2 interacts with host cells and follow up studies have demonstrated how these interactions enhance or diminish virus replication, and/or dampen the innate immune response to infection. The data generated from these interactome studies have also resulted in the identification of possible druggable targets of SARS-CoV-2(11). So far, the proximity dependent proteomic approaches employed in the study of SARS-CoV-2 utilize BioID (19,20) or modified versions of BioID, such as miniTurbo (17) in the study of interactions between SARS-CoV-2 and the host cell. Split-TurboID was developed as a versatile tool for probing the microenvironment of membrane contact sites and the surrounding of protein-protein interactions(21), and has been applied in the profiling of the Ebola virus polymerase interactome (22). However, the approach has not (to our knowledge) been applied in the study of coronavirus-host interactions.

In this study, we fused the inactive N-terminal and C-terminal of split-TurboID to NSP8 and NSP10 of SARS-CoV-2 respectively, and generated A549 cell clones expressing the constructs of interest. As part of the replication and transcription complex of SARS-CoV-2, NSP8 acts as a primase and generates short RNA primers which are required to initiate viral RNA-dependent RNA synthesis. NSP10 on the other hand serves as a cofactor for NSP14 and NSP16, which are involved in proofreading and capping of viral RNA respectively. Our PDL-MS assays resulted in the identification of several proximal partners of SARS-CoV-2 NSP8/ NSP10 complex, a majority of which are RNA-binding proteins. IFN-γ-inducible protein-16 (IFI16), a member of the PYHIN family of proteins, was identified as one of the high-confidence hits. IFI16 has previously been demonstrated to play critical roles in the detection of foreign double stranded DNA in the cytoplasm and the nucleus, as well as restrict the replication of some RNA viruses.Follow-up CRISPR-Cas9 knockout and CRISPRi knockdown of IFI16 however suggested a proviral role for IFI16 during human coronavirus infection.

## Results

### Split-TurboID identifies proximal partners of SARS-CoV-2 NSP8/NSP10 complex

To identify interacting and proximal partners of SARS-CoV-2 non-structural proteins (NSPs), we employed split-TurboID, a versatile tool for proximity labeling in cells (23). The inactive N-terminal or C-terminal of TurboID was fused to either the N-terminal or C-terminal of SARS-CoV-2 NSPs via gateway cloning (Fig. 1a). For controls, FK506-binding protein (FKBP) and the rapamycin-binding domain of mTOR (FRB), two proteins whose association can be induced by rapamycin, were fused to the N and C-terminal fragments of TurboID respectively. To test whether our cloned plasmids can generate functional split-TurboID, pairs of NSPs fused to N- or C-terminal of split-TurboID were transfected into 293T cells and supplemented with 50 uM of biotin. We observed biotinylation of proteins in transfected cells, suggesting the formation of a functional split-TurboID and as expected, there was reduced level of biotinylated proteins in control FKBP/FRB transfected cells that were not supplemented with rapamycin (Fig. S1A). Interestingly, it was observed that fusion of inactive fragments of split-TurboID to the N-or C-terminal of proteins of interest affects the ability to form a functional split-TurboID (Fig. S1A lane 3 and lane 5) and influences overall enzyme efficiency (Fig. S1A lane 5 and lane 9).

**Fig. 1.**
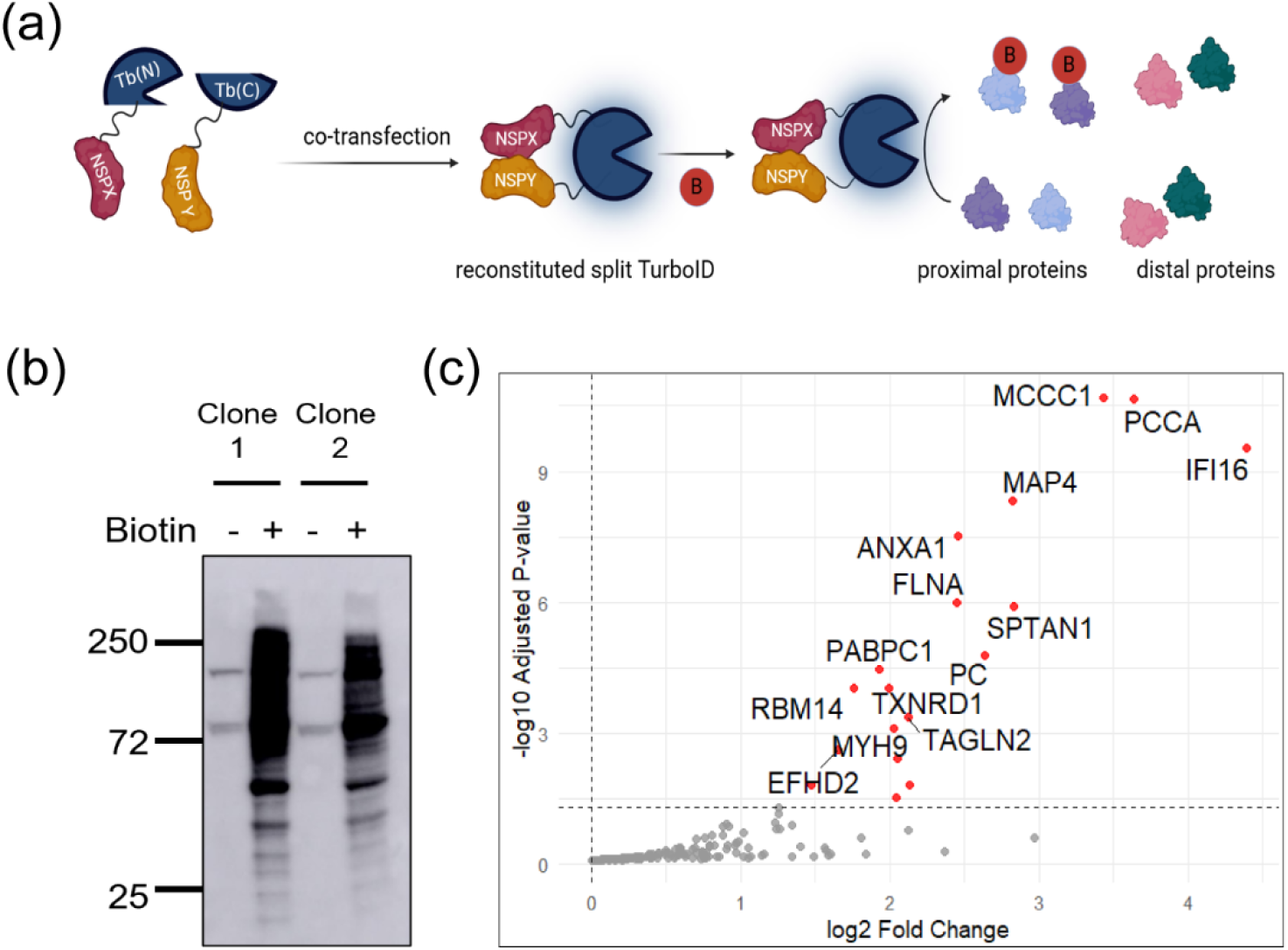
Mass spectrometry identifies proximal partners of SARS-CoV-2 NSP8 and NSP10. (**a)** Schematic representation of how proximity proteomics (split-TurboID) was applied in this study. (**b)** Immunoblot of biotinylated proteins for two clones of A549 cells stably expressing SARS-CoV-2 NSP8-Tb(N) and NSP10-Tb(C). Cells were incubated with or without biotin for 18 h and biotinylated proteins were detected via western blot using streptavidin conjugated to horseradish peroxidase. **(c)** Volcano plot showing enriched host proteins in biotin treated A549 cells (relative to no biotin treated cells) that stably expressed SARS-CoV-2 NSP8-Tb(N) and NSP10-Tb(C).

Based on the results from the transfected 293T cells, we were interested in identifying the host proteins that are likely interacting partners or proximal partners of SARS-CoV-2 NSPs. To achieve this, we generated stable monoclonal A549 cells expressing the NSP/NSP pairs in Fig. S1A via lentivirus transduction. Here, we present data on SARS-CoV-2 NSP8/NSP10 and follow-up experiments, with analyses of additional NSP/NSP pairs to be reported subsequently. Two NSP8/NSP10 cell clones were selected for mass spectrometry experiments, and clones were first screened to confirm expression and formation of a functional split-TurboID (Fig.1b). Biotinylated proteins were isolated by streptavidin pulldown and quantified by liquid chromatography/tandem mass spectrometry. For both cell clones, several proteins were found to be enriched in the biotin treated cells compared to control (untreated) cells. IFI16 was identified as a high confidence proximal partner of SARS-CoV-2 NSP8/NSP10 complex in both cell clones (Fig. 1c and Fig. S1B), and some RNA-binding proteins such as RBM14, PABPC1 and HNRNP A1 were also identified as proximal partners of the complex. A number of carboxylases (PC, PCCA and MCCC1) common to many proteomic studies that employ biotin ligase were also identified, and these are usually considered as background (24,25).

### IFI16 enhances human coronavirus infection

We identified IFI16 as a high-confidence hit in two biological replicates of our mass spectrometry dataset and were curious as to the role of IFI16 in human coronavirus infection. It has earlier been demonstrated that IFI16 restrict Herpesvirus infection (26–29), influenza virus infection (30), and porcine reproductive and respiratory syndrome virus infection (31). To investigate the role of IFI16 in human coronavirus infection, we utilized CRISPR-cas9 to knock out IFI16 in A549 cells and demonstrated loss of IFI16 expression by qRT-PCR and western blot (Fig. 2a, b). To determine the effect of IFI16 loss on coronavirus infection, we infected wildtype (WT) A549 and IFI16-KO cells with Human Coronavirus OC43 (HCoV-OC43) at an MOI of 0.1 for 12 and 24 h. Infected cells were fixed and stained for N protein of HCoV-OC43, followed by immunofluorescence microscopy. We observed a reduced number of nucleocapsid (N)-positive cells in IFI16-KO samples compared to WT at both 12 and 24 h timepoints (Fig. 2c).

**Fig. 2.**
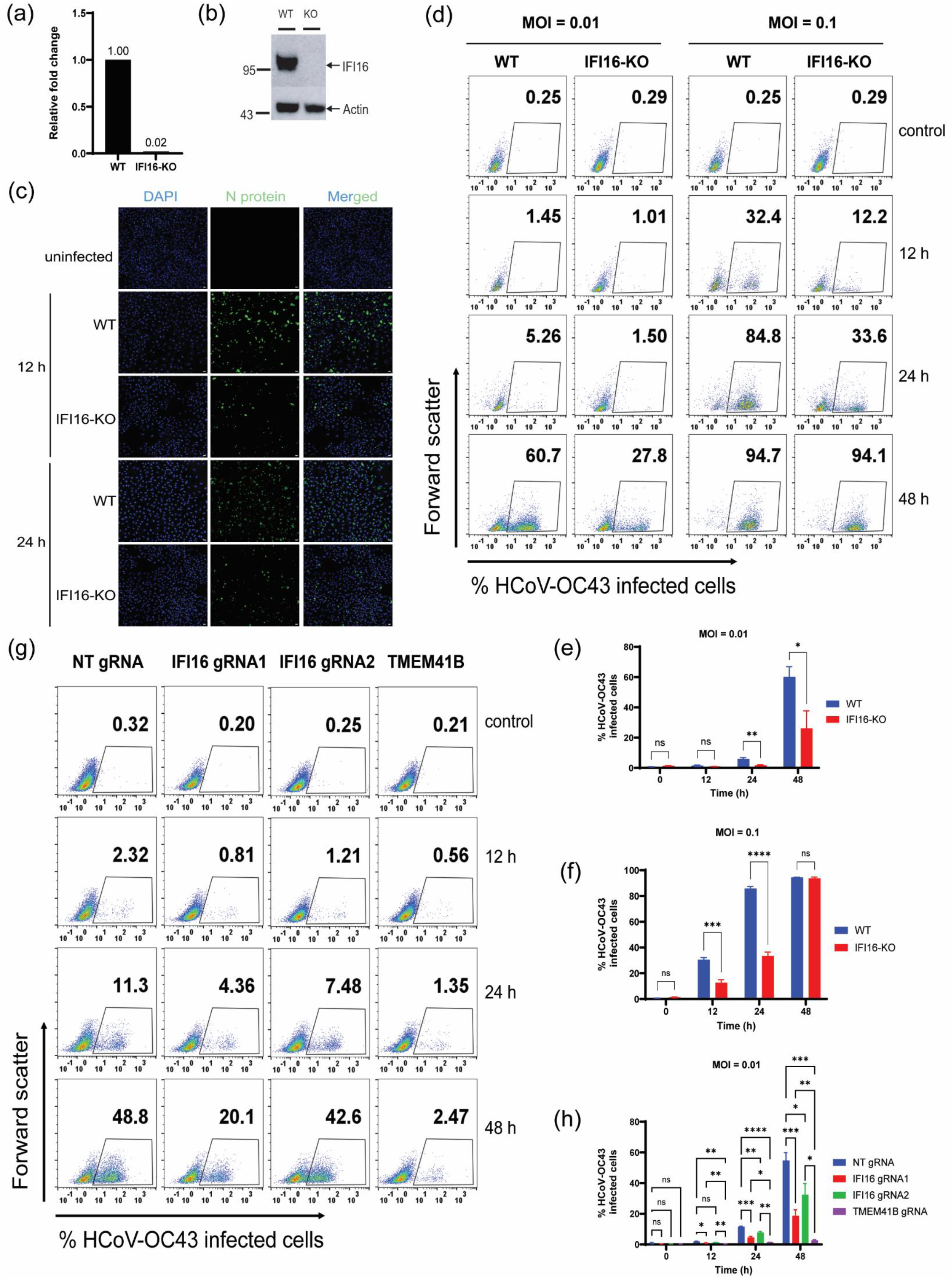
IFI16 promotes coronavirus infection. (**a)** CRISPR-Cas9 technology was used to knock out IFI16 in A549 cells and single cell clones were screened for loss of IFI16 expression using qRT-PCR. Data is representative of two independent experiments. (**b)** Immunoblot showing the loss of IFI16 expression in IFI16-KO cell line. (**c)** Immunofluorescence assay showing reduced number of HCoV-OC43 N-positive cells upon loss of IFI16. Cells were infected at an MOI of 0.1 and incubated for the indicated length of time before fixation and staining for HCoV-OC43 N protein. Scale bar is 10 um. (**d**) Representative flow cytometry analysis of wildtype (WT) A549 cells and IFI16-KO cells infected (in quadruplicate) with HCoV-OC43 at an MOI of 0.01 and 0.1 for 12, 24 and 48 h. Data is representative of 3 independent experiments. **(e)** and **(f)** Quantification of flow cytometry data in Fig. 2d. (**g)** Representative flow cytometry analysis of CRISPRi cell lines infected (in quadruplicate) with HCoV-OC43 at an MOI of 0.01 for 12, 24 and 48 h. data in figure. **(h)** Quantification of flow cytometry data in Fig. 2g. Data is representative of two independent experiments. Statistical significance was assessed by ANOVA. \**P* ≤ 0.05, \*\**P* ≤ 0.01, \*\*\**P* ≤ 0.001, and \*\*\*\**P* ≤ 0.0001.

To assess the impact of IFI16 loss on virus spread, WT and IFI16-KO cells were infected with HCoV-OC43 at an MOI of 0.01. Infected cells were collected, fixed and stained for N protein of HCoV-OC43 at 12, 24 and 48 h post infection. Flow cytometry was then conducted to quantify the number of infected cells at each timepoint, and we observed that the loss of IFI16 results in a decrease in the number of infected cells compared to WT cells at 24 and 48 h post infection (Fig. 2d, e). While an average of 5.8% of WT cells were positive for N protein at 24 h post infection, less than 2% of IFI16-KO cells were positive for viral N protein. At 48 h post infection, an average of 60% of WT cells were positive for N protein compared to 26.1% of IFI16-KO cells (Fig. 2e). These data thus suggests that IFI16 promotes human coronavirus infection, as its absence results in a decrease in infected cells over time.

To evaluate the role of IFI16 in virus replication, we infected wildtype and IFI16-KO cells with HCoV-OC43 at an MOI of 0.1 and conducted flow cytometry to quantify the number of virus infected cells at 12, 24 and 48 h post infection (Fig. 2d, f). While a little over 30% of WT cells were positive for N protein of HCoV-OC43 at 12 h post infection, only 12.6% of IFI16-KO cells were positive for N protein. At 24 h post infection, an average of 85.7% of WT cells were positive for N protein, with only 33.5% of IFI16-KO cells detected as positive for N protein. There was, however, no difference in the number of infected cells between WT and IFI16-KO cells at 48 h post infection. Nonetheless, the difference in virus-infected cells between WT and IFI16-KO cells at 12 and 24 h post infection suggests that IFI16 is likely involved in virus replication and propagation of infection. To determine whether the loss of IFI16 results in increased cell proliferation, which could subsequently affect the number of infected cells observed in our flow cytometry analysis, we compared the proliferation rates of WT and IFI16-KO cells over time and found no significant difference between the two cell lines (Fig. S2C). Previous reports have however demonstrated an antiviral role for IFI16 during DNA and some RNA virus infections (26– 28,30,32). We therefore decided to employ CRISPRi as a secondary method to validate the demonstrated proviral role of IFI16 during coronavirus infection. We designed a non-targeting (NT) gRNA and gRNAs targeting the promoter region of IFI16 and TMEM41B, a previously identified and validated host factor of human coronavirus infection (33).

CRISPRi cell lines expressing gRNAs of interest were infected at an MOI of 0.01 and the number of infected cells that stained positive for N protein was quantified at 12, 24 and 48 h post infection by flow cytometry (Fig. 2g, h and Fig. S2C). At 12 h post infection, an average of 2% of cells expressing NT-gRNA were positive for viral N protein while 0.8% and 1.2% were recorded for cells expressing IFI16-targeting gRNA 1 and 2 respectively (Fig. 2g, h). For cells that express gRNA targeting TMEM41B, an average of 0.5% of cells were positive for N protein. At 24 h post infection, an average of 11.3% of cells expressing NT-gRNA were positive for N protein compared to 4.5% and 7.6% recorded in cells expressing IFI16-gRNA 1 and 2 respectively. Only 1.3% of cells expressing TMEM41B-gRNA were positive for viral N protein. While an average of 54.6% of NT-gRNA expressing cells were positive for N protein of HCoV-OC43 at 48 h post infection, 18.7% and 32.4% of cells expressing IFI16-gRNA 1 and 2 respectively, were positive for viral N protein and only 2.7% of cells expressing TMEM41B-gRNA were positive for viral N protein. The CRISPRi data (Fig. 2h) is thus consistent with earlier CRISPR-KO results (Fig. 2e). The repression of IFI16 expression resulted in a decrease in the number of infected cells compared to cells expressing NT gRNA at all time points post infection, with one gRNA (gRNA1) exhibiting better repression of IFI16. As expected, the repression of TMEM41B, an established host factor of human coronavirus infection resulted in a drastic reduction in the number of infected cells at all time points post infection. Taken together, these data suggest that IFI16 supports human coronavirus infection.

### IFI16 promotes human coronavirus replication

After demonstrating that IFI16 aids HCoV-OC43 infection, we next asked whether IFI16 is involved in viral RNA replication and/or viral protein production. To do this, we conducted time course experiments to investigate the role of IFI16 in coronavirus replication and viral N protein synthesis. WT and IFI16-KO cells were infected at an MOI of 0.1 or 1 and RNA was harvested at different time intervals (4, 8, 12, 16 and 24 h) for qRT-PCR analysis of viral RNA levels. A concurrent experiment was set up, and cells were collected at the same timepoints as RNA was harvested, fixed and stained for viral N protein. Flow cytometry was then conducted to determine the impact of IFI16 loss on viral N protein synthesis. The qRT-PCR analysis revealed that IFI16 plays a role in the early stages of coronavirus replication (Fig. 3a). Upon infection at a low MOI of 0.1, we observed lower levels of viral RNA in IFI16-KO cells compared to WT cells as early as 4 h post infection (Fig. 3a), a time point at which double membrane vesicles (DMVs) have not yet been formed (34). Interestingly, flow cytometry found a concomitant decrease in the number of N-positive cells in IFI16-KO cells compared to WT cells at 4 h post infection (Fig. 3bB). While 20% of WT cells were positive for viral N protein at 4 h post infection, only 2.7% of IFI16-KO cells were positive for N protein. Coronaviruses form DMVs which serve as replication factories within which viral RNA replication occurs, and these are formed within 6 h post infection (34,35) . As such, we observed a significant increase in virus replication in both WT and IFI16-KO cells after 8 h post infection, although viral RNA levels were still lower in IFI16-KO cells compared to WT cells at 8 and 12 h post infection. Interestingly, in WT cells, there was a slight decrease in viral RNA level at 16 h post infection and could be attributed to the release of packaged virions into the cell culture medium. Similar to the trend in viral RNA levels, we observed increasing number of N-positive WT and IFI16-KO cells over time, although the number of infected IFI16-KO cells was lower than that of WT cells at all time points (Fig. 3b and Fig. S3A).

**Fig. 3.**
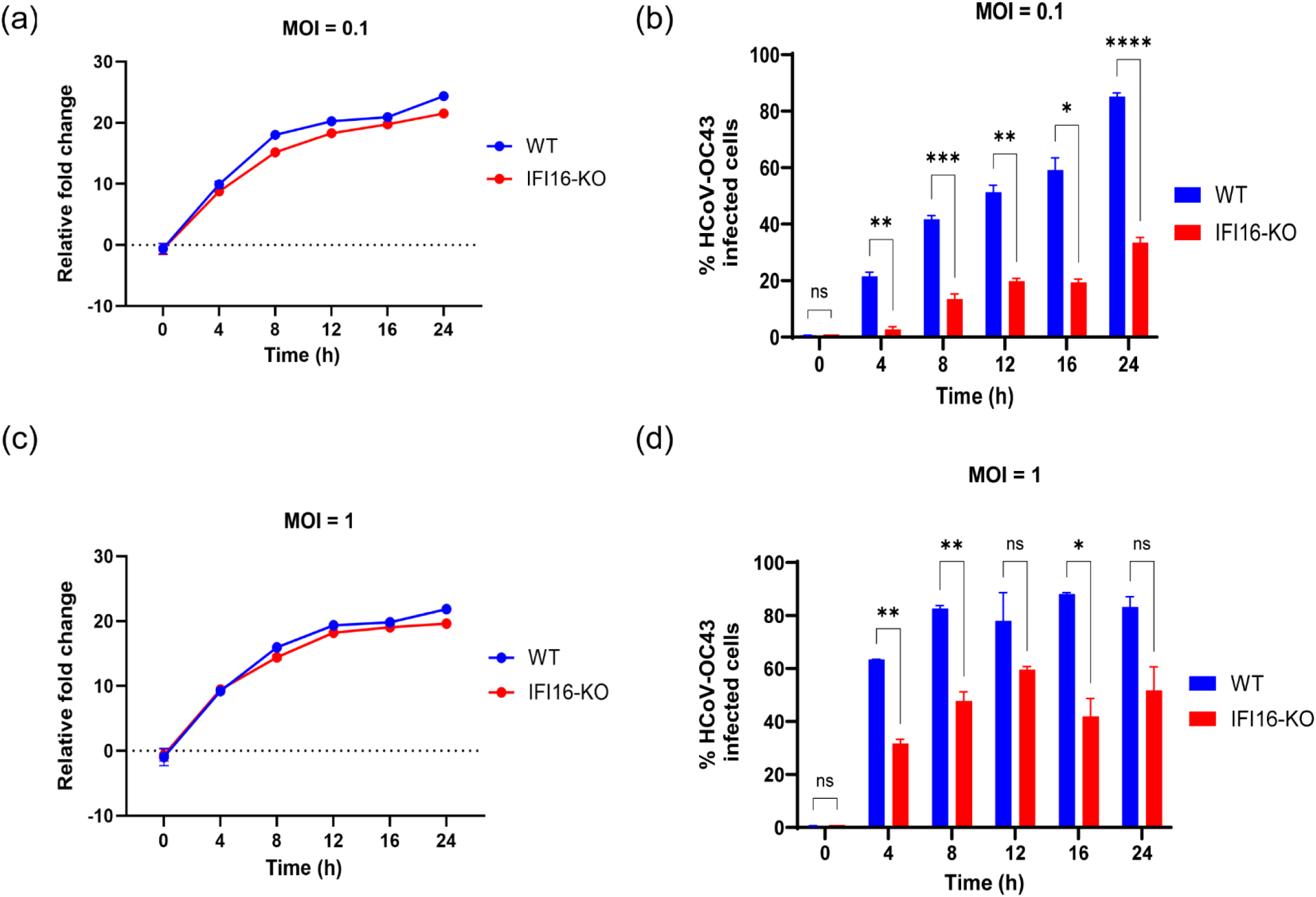
IFI16 enhances coronavirus replication. **(a)** WT and IFI16-KO cells were infected with HCoV-OC43 virus (MOI = 0.1) and RNA was harvested at indicated timepoints post infection for qRT-PCR analysis of N gene levels (n = 2 biologically independent samples). Data is representative of two independent experiments. **(b)** WT and IFI16-KO cells were infected with HCoV-OC43 virus (MOI = 0.1) and cells harvested at indicated timepoints post infection to stain for N protein. Flow cytometry was carried out to quantify the number of virus infected cells at each time point (n = 3 biologically independent samples). Data is representative of two independent experiments. **(c)** WT and IFI16-KO cells were infected with HCoV-OC43 virus (MOI = 1) and RNA was harvested at different times post infection for qRT-PCR analysis of N gene levels (n = 2 biologically independent samples). Data is representative of two independent experiments. **(d)** WT and IFI16-KO cells were infected with HCoV-OC43 virus (MOI = 1) and cells harvested at different timepoints post infection to stain for N protein. Flow cytometry was carried out to quantify the number of virus infected cells at each time point. (n = 4 biologically independent samples). Data is representative of two independent experiments. Statistical significance was assessed by ANOVA. \**P* ≤ 0.05, \*\**P* ≤ 0.01, \*\*\**P* ≤ 0.001, and \*\*\*\**P* ≤ 0.0001.

Upon infection of WT and IFI16-KO cells at a high MOI of 1, we observed no difference in viral RNA levels at 4 h post infection (Fig 3c). However, while 60% of WT cells were positive for viral N protein at 4 h post infection, 31.7% of IFI16-KO cells were positive for viral N protein which represents a relative 50% reduction in number of cells that express viral N protein (Fig. 3d and Fig. S3B). Although viral RNA levels increased in both WT and IFI16-KO cells at 8 and 12 h post infection, there was reduced levels of viral RNA in IFI16-KO cells compared to WT cells (Fig. 3c). A similar trend was observed when we quantified the number of viral N-positive cells at each timepoint (Fig. 3d). At every time point post infection, there was less number of N-positive IFI16-KO cells compared to WT cells. Interestingly, as observed in Fig. 3a, viral RNA levels in WT cells dipped at 16 h post infection and later increased at 24 h post infection (Fig. 3c), and this could be attributed to events stated above. Based on the differences in viral RNA levels between infected IFI16-KO and WT cells, along with the differences in the number of viral N-positive cells over time, we conducted a virus titer assay to determine how the difference in RNA levels affects the production of infectious virus particles. We thus conducted a fluorescent focus forming assay (FFA) to quantify infectious virus particles produced by WT and IFI16-KO cells infected with HCoV-OC43 at an MOI = 0.1 for 12, 24 and 36 h. We demonstrated that the loss of IFI16 results in a decrease in the production of infectious virus particles (Fig. S3C)). At 24 h post infection, the amount of virus produced by infected WT cells was calculated to be approximately 900 FFU/mL, while an estimated 133 FFU/mL was calculated for infected IFI16-KO cells, representing a 6-fold decrease in virus titer.

### The role of IFI16 in coronavirus replication is independent of its role as a transcriptional regulator of type I IFNs

Earlier studies have demonstrated that IFI16 transcriptionally regulates the expression of type I IFNs and other ISGs (36), and others have demonstrated that IFi16 regulates RIG-I expression during IAV infection (30). We thus decided to determine whether the role of IFI16 in coronavirus replication is somehow tied to its role as a transcriptional regulator of type I IFN and RIG-I expression. WT and IFI16-KO cells were infected with HCoV-OC43 and total RNA was harvested at 12 and 24 h post infection. The transcripts of type I IFNs and RIG-I were quantified using qRT-PCR. Compared to WT cells, there was a decrease in the expression of IFNα (Fig. 4a) and IFNβ (Fig. 4b) upon loss of IFI16, suggesting that IFI16 do play a role in the transcriptional regulation of type I IFNs as earlier reported. We also observed a decrease in the expression of RIG-I (Fig. 4c), a gene that is induced by type I IFNs. This suggests that IFI16 promotes human coronavirus infection independent of its role in regulating the transcription of type I IFNs and ISGs. To determine the impact of IFI16 loss on the innate immune response, we transfected WT and IFI16-KO cells with polyinosinic:polycytidylic acid (poly I:C) for 24 h prior to infection with HCoV-OC43 and found no difference in the percentage of coronavirus infected cells for both WT and IFI16-KO cells (Fig. 4c and Fig. S4). This therefore shows that the reduced expression of IFNs upon loss of IFI16 does not affect the ability of cells to mount a robust innate immune response to virus infection. Taken together, these data suggests that although the loss of IFI16 results in reduced production of type I IFNs during HCoV-OC43 infection, its role in aiding coronavirus replication appear to outweigh its role as a transcriptional regulator of type I IFN expression (Fig. 4c, Fig. S4).

**Fig. 4.**
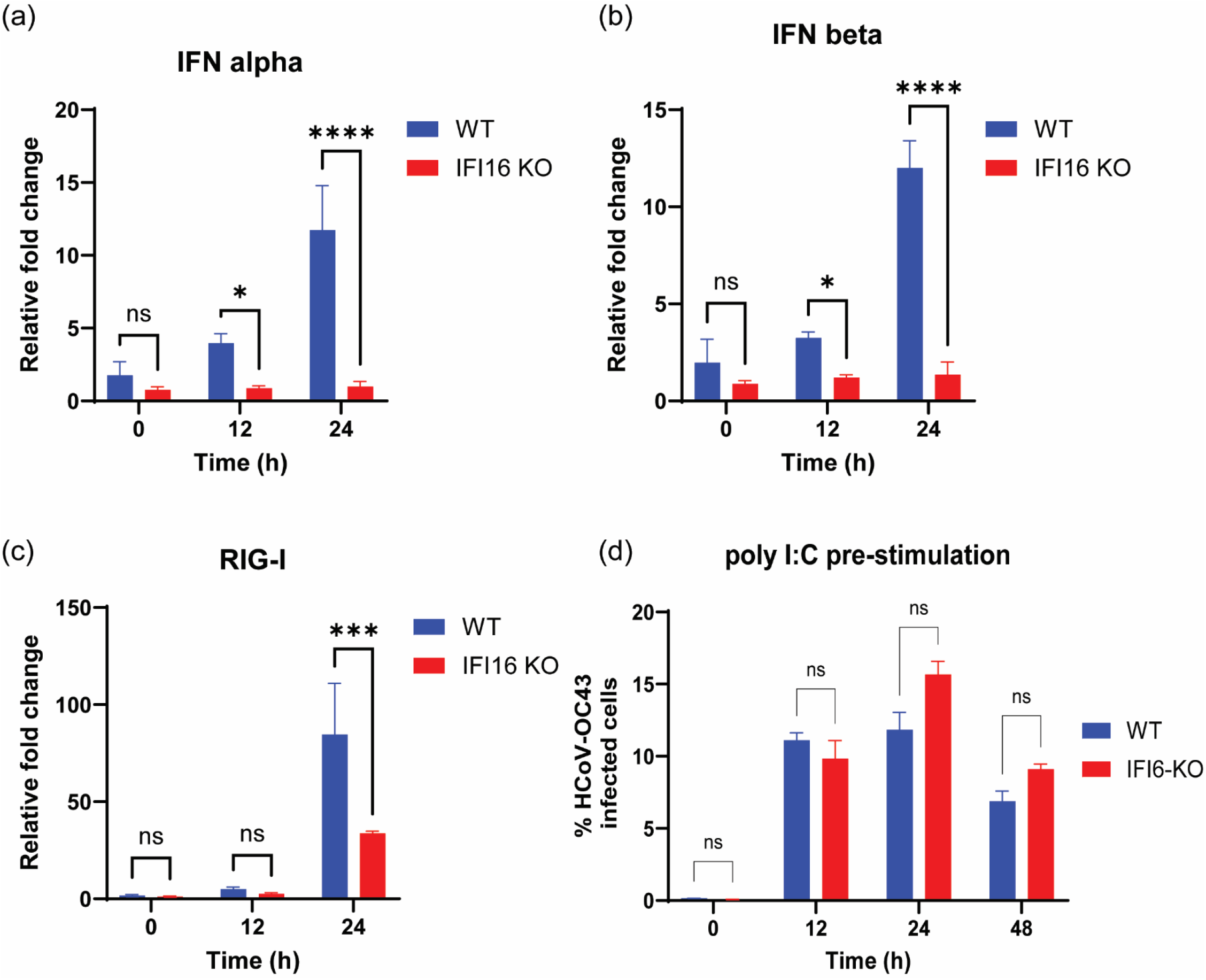
The role of IFI16 in coronavirus replication is independent of its role as a transcriptional regulator of type I IFN genes. WT and IFI16-KO cells were infected with HCoV-OC43 virus (MOI = 0.1) and RNA was harvested at 12 and 24hr post infection for qRT-PCR analysis of the expression levels of type I IFNs (**a** and **b**) and RIG-I (**c**). Data is representative of two independent experiments (n = 3 biologically independent samples). **(d)** WT and IFI16-KO cells were stimulated with poly I:C for 24 h and then infected with HCoV-OC43 virus (MOI = 1) for 12, 24 and 48 h. Cells were harvested and stained for N protein, followed by flow cytometry. Statistical significance was assessed by ANOVA. \**P* ≤ 0.05, \*\*\**P* ≤ 0.001, and \*\*\*\**P* ≤ 0.0001

## Discussion

We utilized proximity proteomics to determine host proteins that are proximal or potential interacting partners of coronavirus NSP8 and NSP10 and identified several proteins that are bonafide RNA-binding proteins (Fig.1b, Fig. S1B ), some of which have previously been identified in interactome studies involving live virus infection (37–40). Among the list of identified proteins include ANXA1, HNRNPA1, RBM14, RNF213, PABPC1, and IFI16. Annexin A1 (ANXA1) is a phospholipid-binding, calcium-dependent protein that play essential roles in a variety of cellular functions including inflammation, proliferation, and apoptosis (41). Heterogeneous nuclear ribonucleoprotein A1 (HNRNP A1) belongs to the HNRNP family of proteins (42), and has been shown to interact with the nucleocapsid (N) protein of Mouse hepatitis virus (MHV) and SARS-CoV to regulate viral RNA synthesis (43–45). RNA binding motif 14 (RBM14) is an RNA-binding protein that has been demonstrated to regulate the replication of DNA and RNA viruses (46–48). Ring Finger Protein 213 (RNF213) is a large protein that has been found to possess ubiquitin ligase and ATPase activities (49,50), and serves as a sensor of ISGylated proteins that are associated with lipid droplets (51). Knockdown of RNF213 resulted in increased HSV-1 and Coxsackievirus B3 (CVB3) infection in HeLa cells, as well as an increase in Respiratory syncytial virus (RSV) infection in A549 cells (51). Polyadenylate-binding protein cytoplasmic 1 (PABPC1) is an RNA-binding protein that is involved in mRNA translation, mRNA localization and mRNA decay (52). Previous studies have shown that PABPC1 promotes PEDV replication by enhancing the translation of viral mRNA during infection in Vero cells (53,54), and has recently been demonstrated to be involved in the capping of SARS-CoV-2 viral mRNA (55). Interestingly, PABPC1 along with stress granule proteins were identified in SARS-CoV-2 virions (40).

IFN-γ-inducible protein-16 (IFI16), an interferon stimulated gene (ISG), was identified in this study as a high-confidence hit in two separate proximity proteomics experiments. IFI16 is a member of the pyrin and HIN domain (PYHIN) containing protein family, which consist of proteins that are involved in the regulation of the innate immune response to foreign DNA (56,57). While the pyrin domain of IFI16 is involved in protein-protein interactions (58), the HIN domains promote DNA-binding in a non-sequence specific manner, which allows for the detection of diverse microbial DNA (27,59,60). The localization of IFI16 is cell type-dependent (61), and it has been demonstrated that the acetylation of a nuclear localization signal at the N-terminal region of IFI16 regulates its cellular distribution (62).

Initial studies reported that IFI16 acts as a transcriptional repressor (63). Subsequent studies however demonstrated that IFI16 promotes the transcription of type I IFNs and IFN-stimulated genes to enhance IFN response to a variety of innate immune stimuli (36). Other studies have shown that IFI16 acts as an innate immune sensor for intracellular DNA (64), and can restrict the replication of several nucleus-replicating viruses such as human cytomegalovirus (HCMV) (26), herpes simplex virus (HSV) (27), and human papillomavirus 18 (HPV18) (28). IFI16 has also been demonstrated to act as a cytosolic immune sensor of HIV-1 DNA species in macrophages (65), by initiating an IFN response via the cyclic GMP-AMP synthase (cGAS) and stimulator of interferon genes (STING) pathway (66). Aside from the detection and restriction of DNA viruses, IFI16 is reported to be involved in the detection of HIV-1 reverse transcription (RT) intermediates, which results in caspase-1 dependent pyroptotic death of abortively infected CD^4+^ T cells (67).

Using CRISPR-Cas9 knock out and CRISPRi studies, we have demonstrated that the loss of IFI16 results in a reduced coronavirus replication, thus suggesting that IFI16 promotes human coronavirus infection. Previous studies, however, identified an antiviral role for IFI16 in the context of both positive and negative-sense single-stranded RNA viruses. Overexpression of IFI16 was reported to restrict the replication of Chikungunya virus (CHIKV) and Zika virus (ZIKV) (32), two positive-sense single-stranded RNA viruses that are transmitted by mosquitoes. *In vitro* and *in vivo* studies have also found that IFI16inhibits the replication of IAV, a segmented negative-sense RNA virus (30). Mechanistically, IFI16 was found to directly bind IAV RNA to augment RIG-I activation and gene transcription. While others have demonstrated an antiviral role for IFI16 against other positive-sense single-stranded RNA viruses (32), we have demonstrated a proviral role of IFI16 during human coronavirus infection, thus suggesting a virus-specific function of IFI16.

Comparative analysis of various studies reveals a dichotomy of function by host proteins during virus infection. Depending on the virus that infects a cell, some host proteins either play antiviral roles or are hijacked by viruses to aid in virus replication. For example, HNRNP A1 has been demonstrated to interact with the N protein of IAV which results in restriction of IAV replication (68), but the interaction of HNRNP A1 with the N protein of MHV and SARS-CoV-2 promotes virus replication (44,45,69). Also, while RBM14 has been found to inhibit replication of Porcine epidemic diarrhea virus (PEDV) by upregulating the expression of type I IFN (48), the NS1 Protein of IAV has been shown to promote relocalization of RBM14 to the nucleolus to promote IAV replication (46). The differential activities of hnRNPA1 and RBM14 against positive sense (coronavirus) and negative sense (IAV) RNA viruses highlights the virus-specific role of some host proteins during infection.

Interferon stimulated genes (ISGs) are gene products of the interferon (IFN)-mediated innate immune response that play diverse roles in the control of pathogens (70), and several ISGs play antiviral roles by targeting various stages of a virus life cycle. Although a majority of identified ISGs have been found to possess antiviral activity against a broad spectrum of DNA and RNA viruses (71), recent studies have demonstrated the possibility of ISGs to be repurposed by viruses to support virus replication. IFIT2 (interferon-induced protein with tetratricopeptide repeats 2), a member of the IFIT family of proteins which are known to exert antiviral activity against a diverse number of DNA and RNA viruses, was demonstrated to promote translation of IAV mRNAs (72). IFIT2 was found to associate with active ribosomes and this interaction inhibited ribosome pausing on viral mRNAs, thus resulting in efficient viral protein translation. IFIT3 (interferon-induced protein with tetratricopeptide repeats 3), also a member of the IFIT family of proteins, was also found to promote IAV infection, by enhancing the translational efficiency of viral proteins (73). Using two independent CRISPR-cas9 approaches, we have demonstrated that IFI16, an ISG, promotes HCoV-OC43 replication (Fig. 2) and its loss results in reduced virus replication and production of infectious virus particles (Fig. 3, Fig. S3C).

Previous studies have demonstrated a role of IFI16 in the transcriptional regulation of type I IFNs genes, as well as some interferon stimulated genes (36). Compared to WT cells, we observed reduced expression of type I IFNs in IFI16-KO cells (Fig. 4), thus supporting the role of IFI16 in regulating the expression of type I IFNs and other interferon stimulated genes. Others have also demonstrated a role of IFI16 in the expression of RIG-I (30), and we have shown that the loss of IFI16 results in reduced expression of RIG-I during virus infection. Taken together, these data suggests that the role of IFI16 in human coronavirus infection is independent of its role as a transcriptional regulator of type I IFN gene expression.

## Methods

### Plasmids

Gateway cloning was utilized to clone Plasmids encoding the non-structural proteins of SARS-CoV-2 fused to inactive N-terminal or C-terminal fragments of split TurboID were cloned into pLEX_307 (Addgene: 41392) via Gateway cloning, and all plasmids were sequenced to confirm the integrity of each fusion construct. Control plasmids (FKBP12 and FRB fused to N-terminal and C-terminal of split-TurboID respectively) were also cloned via Gateway cloning.

### Cells and Virus infections

HEK 293T and A549 cells were maintained in Dulbecco’s modified Eagle’s medium (DMEM) (Thermo Fisher) supplemented with 10% bovine calf serum (BCS) (Thermo Fisher) and 1% penicillin-streptomycin (P/S) (Thermo Fisher). Virus infection of A549 cells was performed using DMEM supplemented with 2% BCS and 1% P/S). HCT-8 cells were maintained in Roswell Park Memorial Institute Medium (RPMI) supplemented with 10% BCSand 1% P/S. Human Coronavirus-OC43 was obtained from Biodefense and Emerging Infections Research Resources Repository (BEI Resources) and propagated in HCT-8 cells supplemented with no serum and 1 (P/S). The virus was passaged 3 more times and passage four was used for infection assays.

For flow cytometry assays, 1 x 10^5^ cells were seeded in 24-well plates and incubated overnight. The next day, cells were washed once with 1X PBS and 100 uL of infection media (DMEM, 2% FCS, 1% P/S) was placed in each well followed by the addition of diluted virus stock to obtain the desired MOI and plates incubated at 33 ^°^C for 1 h and 30 min, with periodic swirling of plates every 20 min. Media was replaced with 500 uL of fresh infection media and plates incubated at 33 ^°^C. Infected cells were harvested at the indicated timepoints post infection and fixed using 4% PFA for 10 min at 4°C. Cells were washed twice with a permeabilization/wash buffer (0.1% saponin and 0.5% BSA in 1X PBS) and stained with a 1:10000 dilution of anti-HCoV-OC43-N rabbit polyclonal antibody (catalogue no. 40643-T62; Sino Biological) for 30 min at 4°C. Cells were washed twice and incubated with a 1:1000 dilution of goat anti-rabbit antibody conjugated to Alexa Fluor 488 (catalogue no. 4030-30; SouthernBiotech) for 30 min at 4°C. Cells were washed twice after staining, resuspended in 1X PBS containing 1% BCS and flow cytometry was carried out to quantify the number of infected cells at each time point. For poly I:C prestimulation of cells, 1 x 10^5^ cells were seeded in 24-well plates overnight, followed by transfection of 200 ng of poly I:C for 24 h using X-tremeGENE (Roche). Cells were then infected at an MOI = 1 and at the indicated time points post infection, infected cells were collected and stained for N protein of HCoV-OC43. Flow cytometry was then carried out to quantify the number of infected cells at each timepoint.

For fluorescence imaging assays, 1 x 10^5^ cells were seeded on cover slips in 24-well plates and incubated overnight, followed by infection with HCov-OC43 for the indicated time points. Cells were fixed with 4% PFA for 10 min at room temperature and washed three times with PBS. (2% BSA/PBST) for 1 h and incubated with primary antibody (anti-HCoV-OC43-N rabbit polyclonal antibody) overnight at 4°C. Cells were washed three times with PBST and incubated with secondary antibody (goat anti-rabbit antibody conjugated to Alexa Fluor 488) and DAPI for 1 h at room temperature. Cells were washed three times in PBST and mounted for immunofluorescence microscopy.

### Mass spectrometry

Samples for mass spectrometry were processed as earlier described (74), with some modifications. About 15 x 10^6^ A549 cells stably expressing SARS-CoV-2 NSP8-Tb(N) and NSP10-Tb(C) were seeded in T175 flasks cells for 24 h, after which cells were left untreated or treated with 50 μM of biotin and incubated for 18 h. Cells were trypsinized and collected by centrifugation at 500 g for 5 min at 4°C. The cells were then lysed by resuspension in RIPA lysis buffer containing 1X protease inhibitor (Catalog number A32955) and incubated for 10 min at 4°C. Lysate clarified by centrifuging at 12,000 g for 10 min at 4°C and protein concentration was determined using BCA protein assay kit (Thermo Fisher). A 25 uL volume of streptavidin magnetic beads was placed in 1.7 mL Eppendorf tubes and washed twice with 1 mL of RIPA lysis buffer (using a magnet to separate beads from the lysis buffer each time). Beads were then incubated with 300 ug of protein from biotin treated and untreated sample and an additional 500 μL of RIPA lysis buffer containing 1X protease inhibitor was added. The mixture was incubated at 4 °C with rotation overnight.

To prepare samples for mass spectrometry analysis, proteins bound to streptavidin beads were washed with 200 μL of 50 mM Tris-HCl (pH 7.5), followed by two washes with 200 μL 2M urea in 50 mM Tris (pH 7.5) buffer. On-bead trypsin digestion was then carried out by incubating beads with 80 μL of 2 M urea in 50 mM Tris-HCL containing 1 mM Dithiothreitol and 0.4 μg trypsin at 25 °C for 1 h while shaking at 1,000 r.p.m. The supernatant was transferred to fresh tubes and beads were washed twice with 60 μL of 2 M urea in 50 mM Tris (pH 7.5) buffer and the washes were combined with the on-bead digested supernatant.

Samples were further processed using the single-pot solid-phase-enhanced sample preparation technology (75), with some modifications. Briefly, to reduce and alkylate digested proteins, 7.5 uL of Tris-(2-Carboxyethyl)phosphine, Hydrochloride (200 mM) was added to samples and incubated at 25 °C with shaking (12 000 r.p.m) for 20 mins, followed by the addition of 7.25 uL of Iodoacetic Acid (500 mM) and further incubation for another 20 mins. A 1:1 mix of Sera-Mag SpeedBeads (GE Healthcare, cat. no. 45152105050250) and (GE Healthcare, cat. no. 65152105050250) was prepared and washed 3 times with distilled water and 10 uL bead preparation was added to each sample, followed by the addition of 350 uL of 50% ethanol and rotation of samples at room temperature for 10 min. Beads were magnetized and washed 3 times with 500 uL of 80% ethanol. Beads were allowed to dry after the final wash and 30uL of Triethylammonium bicarbonate buffer (50 mM) was added to each sample, followed by the addition of mass spectrometry grade lys-C and trypsin. Samples were then incubated overnight at 37 °C with shaking at 12 000 r.p.m overnight. To clean up peptides, 1 mL of Acetonitrile (ACN) was added to samples to bring ACN concentration to > 95% and samples rotated at room temperature for 10 mins. Beads were magnetized and washed 3 times with 500 uL of pure ACN, followed by elution of peptides using 2% Dimethylsulfoxide. A speed vacuum was used to dry down the samples and samples resuspended in 10 uL of 5% formic acid for analysis by liquid chromatography-tandem mass spectrometry (LC-MS/MS).

Peptides were separated on a 75 µm × 25 cm in-house packed C18 column coupled to a Dionex Ultimate 3000 nanoflow UHPLC system. A 140-min gradient of increasing acetonitrile (ACN) was applied at a flow rate of 200 nL/min. Mass spectrometric data were acquired on an Orbitrap Fusion Lumos Tribrid mass spectrometer operating in data-dependent acquisition (DDA) mode. Full MS scans were acquired in the Orbitrap at 120,000 resolution, with an automatic gain control (AGC) target of 2 × 10^5^ and a maximum injection time of 100 ms. MS/MS spectra were acquired at 15,000 resolution following precursor isolation with a 1.6 m/z isolation window and higher-energy collisional dissociation (HCD) using 35% normalized collision energy. A 3 s cycle time was employed to select precursors for MS/MS fragmentation from each full MS scan. Dynamic exclusion was enabled with an exclusion duration of 25 s. MaxQuant was used to analyze raw mass spectrometry data, and further analysis was carried out using R (Version 4.5.1; R Core Team, 2025) and RStudio (Version 2025.09.2+418; Posit Team, 2025).

### Western Blot

To identify NSP-NSP interactions that result in the formation of a functional split-TurboID, HEK 293T cells were seeded in 6-well plates and transfected with NSP pairs (fusion constructs) for 24 h, after which media was replaced with fresh complete medium containing 50 uM of biotin. For cells transfected with FKBP12 and FRB, rapamycin was added (positive control) or not added (negative control) during media replacement and cells were incubated for 18 h. Cells were collected by centrifugation at 500 g for 5 min at 4°C and lysed using RIPA lysis buffer containing 1X protease inhibitor. Cell lysates were clarified by centrifugation at 12,000 g for 10 min at 4°C and 30 μg of whole-cell lysate was combined with 2X Laemmli buffer and samples boiled at 95 °C for 10 min. Proteins were separated by SDS-PAGE and transferred onto a nitrocellulose membrane, and membrane was blocked with 5%(w/v) non-fat milk in 1X TBST at room temperature for 30 min. Biotinylated proteins were detected by incubating membrane with 0.3 μg/mL streptavidin–HRP in 3% (w/v) BSA in 1X TBST at 4°C overnight. Membrane was washed three times with 1X TBST and developed using Pierce™ ECL western blotting substrate (Thermo Fisher). For the screening of IFI16-KO cell clones, cells were lysed using RIPA lysis buffer containing 1X protease inhibitor and cell lysates were clarified by centrifugation at 12, 000 g for 10 min at 4°C. Protein concentration was determined using the BCA protein assay kit (Thermo Fischer) and 30 μg of whole-cell lysate was combined with 2X Laemmli buffer and samples boiled at 95 °C for 10 min. Proteins were separated by SDS-PAGE and transferred onto a nitrocellulose membrane, and membrane was blocked with 5%(w/v) non-fat milk in 1X TBST at room temperature for 30 min. Membrane was probed for IFI16 using anti-IFI16 antibody from Cell Signaling Technology (mAb: 14970).

### CRISPR-KO and CRISPRi

CRISPR-Cas9 was utilized to knock out IFI16 in A549 cells. Briefly, sgRNA targeting exon 2 of IFI16 was designed using Synthego software and supplied by Synthego. The sgRNA and Cas9 mRNA were electroporated into A549 cells using the Neon electroporation system (Invitrogen) and cells plated in antibiotic free media to recover. Cells were expanded and single cell cloning was carried out, with clones screened for loss of IFI16 by qRT-PCR and western blot. For the generation of CRISPRi cell lines, A549 cells were transduced with a lentivirus preparation of dCAS9-KRAB-mCherry for 48 h, after which cells were sorted for mCherry positive cells and expanded. Non-targeting sgRNA and sgRNAs targeting the promoter regions of IFI16 or TMEM41B were designed using Benchling software, cloned into lentiGuide-Puro plasmid (Addgene: 52963) and sanger sequenced for confirmation of gRNA sequences. Sorted A549 cells expressing dCas9-KRAB were expanded and transduced with lentivirus preparation of non-targeting sgRNA (negative control) or sgRNAs targeting the promoter region of IFI16 or TMEM41B (positive control). Transduced cells were incubated for 48 h, after which puromycin was added to select for successfully transduced cells. Puromycin selected cells were then expanded for use in experiments. c

### Quantitative real-time PCR (qRT-PCR)

Cells were seeded in 24-well plates overnight and infected with HCoV-OC43 at the indicated multiplicity of infection (MOI). Total RNA was harvested and quantified using a NanoDrop (Thermo Scientific). A total of 300 ng of RNA was used to generate cDNA using protoscript II reverse transcriptase (M0368, New England biolabs Inc) and diluted 1:20 in DNAse free water before being used as template for qPCR assays. The Taqman assay for IFI16 (Applied Biosystems, ID: Hs00986757_m1) was employed in the detection of IFI16 expression following the manufacturers protocol. For the detection of N gene and type I IFNs, qRT-PCR assays were performed using the Luna Universal Probe qPCR Master Mix (M3003, New England biolabs Inc) on QuantStudio 3. The Ct values from the Design and Analysis software were exported to excel for calculation of fold changes and data was imported into GraphPad for statistical analysis. Primers used in this study are listed on Table 1.

## Acknowledgements

We thank Prof. Christopher Huston (University of Vermont) for generously providing HCT-8 cells for this work.

## Abbreviations

NSP: non-structural protein
IFI16: IFN-γ-inducible protein-16
HCoV-OC43: Human coronavirus OC43
CRISPR: Clustered Regularly Interspaced Short Palindromic Repeats
CRISPRi: Clustered Regularly Interspaced Short Palindromic Repeats Interference
MOI: multiplicity of infection.

## Author contributions

D.M. and S.L designed the research. S.L. conducted the experiments. I.B., Z.M., M.S., J.S., J.D., W.D., T.K., and J.W. provided technical support for the experiments. S.L and D.M. analyzed the data and wrote the paper.

## References

1. Corman VM, Muth D, Niemeyer D, Drosten C. Chapter Eight - Hosts and Sources of Endemic Human Coronaviruses. In: Kielian M, Mettenleiter TC, Roossinck MJ, editors. Advances in Virus Research [Internet]. Academic Press; 2018 [cited 2025 Jan 5]. p. 163–88. Available from: https://www.sciencedirect.com/science/article/pii/S0065352718300010

2. Coronaviridae Study Group of the International Committee on Taxonomy of Viruses, Gorbalenya AE, Baker SC, Baric RS, De Groot RJ, Drosten C, et al. The species Severe acute respiratory syndrome-related coronavirus: classifying 2019-nCoV and naming it SARS-CoV-2. Nat Microbiol. 2020 Mar 2;5(4):536–44.

3. Gorbalenya AE, Enjuanes L, Ziebuhr J, Snijder EJ. Nidovirales: Evolving the largest RNA virus genome. Virus Res. 2006 Apr;117(1):17–37.

4. Kesheh MM, Hosseini P, Soltani S, Zandi M. An overview on the seven pathogenic human coronaviruses. Rev Med Virol. 2022;32(2):e2282.

5. Corman VM, Muth D, Niemeyer D, Drosten C. Hosts and Sources of Endemic Human Coronaviruses. Adv Virus Res. 2018;100:163–88.

6. Cimolai N. Complicating Infections Associated with Common Endemic Human Respiratory Coronaviruses. Health Secur. 2021 Apr;19(2):195–208.

7. Cui J, Li F, Shi ZL. Origin and evolution of pathogenic coronaviruses. Nat Rev Microbiol. 2019 Mar;17(3):181–92.

8. Steiner S, Kratzel A, Barut GT, Lang RM, Aguiar Moreira E, Thomann L, et al. SARS-CoV-2 biology and host interactions. Nat Rev Microbiol. 2024 Apr;22(4):206–25.

9. Davies JP, Almasy KM, McDonald EF, Plate L. Comparative Multiplexed Interactomics of SARS-CoV-2 and Homologous Coronavirus Nonstructural Proteins Identifies Unique and Shared Host-Cell Dependencies. ACS Infect Dis. 2020 Dec 11;6(12):3174–89.

10. Gordon DE, Hiatt J, Bouhaddou M, Rezelj VV, Ulferts S, Braberg H, et al. Comparative host-coronavirus protein interaction networks reveal pan-viral disease mechanisms. Science. 2020 Dec 4;370(6521):eabe9403.

11. Gordon DE, Jang GM, Bouhaddou M, Xu J, Obernier K, White KM, et al. A SARS-CoV-2 protein interaction map reveals targets for drug repurposing. Nature. 2020 Jul;583(7816):459–68.

12. Li J, Guo M, Tian X, Wang X, Yang X, Wu P, et al. Virus-Host Interactome and Proteomic Survey Reveal Potential Virulence Factors Influencing SARS-CoV-2 Pathogenesis. Med. 2021 Jan 15;2(1):99–112.e7.

13. Zhou Y, Liu Y, Gupta S, Paramo MI, Hou Y, Mao C, et al. A comprehensive SARS-CoV-2–human protein– protein interactome reveals COVID-19 pathobiology and potential host therapeutic targets. Nat Biotechnol. 2022 Oct 10;1–12.

14. Graziadei A, Schildhauer F, Spahn C, Kraushar M, Rappsilber J. SARS-CoV-2 Nsp1 N-terminal and linker regions as a platform for host translational shutoff [Internet]. bioRxiv; 2022 [cited 2025 Jan 5]. p. 2022.02.10.479924. Available from: https://www.biorxiv.org/content/10.1101/2022.02.10.479924v2

15. Burnap SA, Ortega-Prieto AM, Jimenez-Guardeño JM, Ali H, Takov K, Fish M, et al. Cross-Linking Mass Spectrometry Uncovers Interactions Between High-Density Lipoproteins and the SARS-CoV-2 Spike Glycoprotein. Mol Cell Proteomics [Internet]. 2023 Aug 1 [cited 2025 Jan 5];22(8). Available from: https://www.mcponline.org/article/S1535-9476(23)00111-1/abstract

16. Laurent EMN, Sofianatos Y, Komarova A, Gimeno JP, Tehrani PS, Kim DK, et al. Global BioID-based SARS-CoV-2 proteins proximal interactome unveils novel ties between viral polypeptides and host factors involved in multiple COVID19-associated mechanisms [Internet]. bioRxiv; 2020 [cited 2025 Jan 5]. p. 2020.08.28.272955. Available from: https://www.biorxiv.org/content/10.1101/2020.08.28.272955v1

17. Samavarchi-Tehrani P, Abdouni H, Knight JDR, Astori A, Samson R, Lin ZY, et al. A SARS-CoV-2 – host proximity interactome [Internet]. bioRxiv; 2020 [cited 2024 Oct 20]. p. 2020.09.03.282103. Available from: https://www.biorxiv.org/content/10.1101/2020.09.03.282103v1

18. St-Germain JR, Astori A, Samavarchi-Tehrani P, Abdouni H, Macwan V, Kim DK, et al. A SARS-CoV-2 BioID-based virus-host membrane protein interactome and virus peptide compendium: new proteomics resources for COVID-19 research [Internet]. bioRxiv; 2020 [cited 2025 Jan 5]. p. 2020.08.28.269175. Available from: https://www.biorxiv.org/content/10.1101/2020.08.28.269175v1

19. Meyers JM, Ramanathan M, Shanderson RL, Beck A, Donohue L, Ferguson I, et al. The proximal proteome of 17 SARS-CoV-2 proteins links to disrupted antiviral signaling and host translation. PLOS Pathog. 2021 Oct 1;17(10):e1009412.

20. May DG, Martin-Sancho L, Anschau V, Liu S, Chrisopulos RJ, Scott KL, et al. A BioID-Derived Proximity Interactome for SARS-CoV-2 Proteins. Viruses. 2022 Mar;14(3):611.

21. Cho KF, Branon TC, Rajeev S, Svinkina T, Udeshi ND, Thoudam T, et al. Split-TurboID enables contact-dependent proximity labeling in cells. Proc Natl Acad Sci. 2020 Jun 2;117(22):12143–54.

22. Fang J, Pietzsch C, Tsaprailis G, Crynen G, Cho KF, Ting AY, et al. Functional interactomes of the Ebola virus polymerase identified by proximity proteomics in the context of viral replication. Cell Rep. 2022 Mar 22;38(12):110544.

23. Cho KF, Branon TC, Rajeev S, Svinkina T, Udeshi ND, Thoudam T, et al. Split-TurboID enables contact-dependent proximity labeling in cells. Proc Natl Acad Sci. 2020 Jun 2;117(22):12143–54.

24. Kim DI, Jensen SC, Roux KJ. Identifying Protein-Protein Associations at the Nuclear Envelope with BioID. Methods Mol Biol Clifton NJ. 2016;1411:133–46.

25. Roux KJ, Kim DI, Burke B, May DG. BioID: A Screen for Protein-Protein Interactions. Curr Protoc Protein Sci. 2018 Feb 1;91:19.23.1-19.23.15.

26. Gariano GR, Dell’Oste V, Bronzini M, Gatti D, Luganini A, Andrea MD, et al. The Intracellular DNA Sensor IFI16 Gene Acts as Restriction Factor for Human Cytomegalovirus Replication. PLOS Pathog. 2012 Jan 26;8(1):e1002498.

27. Johnson KE, Bottero V, Flaherty S, Dutta S, Singh VV, Chandran B. IFI16 Restricts HSV-1 Replication by Accumulating on the HSV-1 Genome, Repressing HSV-1 Gene Expression, and Directly or Indirectly Modulating Histone Modifications. PLOS Pathog. 2014 Nov 6;10(11):e1004503.

28. Lo Cigno I, De Andrea M, Borgogna C, Albertini S, Landini MM, Peretti A, et al. The Nuclear DNA Sensor IFI16 Acts as a Restriction Factor for Human Papillomavirus Replication through Epigenetic Modifications of the Viral Promoters. J Virol. 2015 Jul 8;89(15):7506–20.

29. Roy A, Dutta D, Iqbal J, Pisano G, Gjyshi O, Ansari MA, et al. Nuclear Innate Immune DNA Sensor IFI16 Is Degraded during Lytic Reactivation of Kaposi’s Sarcoma-Associated Herpesvirus (KSHV): Role of IFI16 in Maintenance of KSHV Latency. J Virol. 2016 Sep 12;90(19):8822–41.

30. Jiang Z, Wei F, Zhang Y, Wang T, Gao W, Yu S, et al. IFI16 directly senses viral RNA and enhances RIG-I transcription and activation to restrict influenza virus infection. Nat Microbiol. 2021 Jul;6(7):932–45.

31. Chang X, Shi X, Zhang X, Wang L, Li X, Wang A, et al. IFI16 Inhibits Porcine Reproductive and Respiratory Syndrome Virus 2 Replication in a MAVS-Dependent Manner in MARC-145 Cells. Viruses. 2019 Dec 16;11(12):1160.

32. Wichit S, Hamel R, Yainoy S, Gumpangseth N, Panich S, Phuadraksa T, et al. Interferon-inducible protein (IFI) 16 regulates Chikungunya and Zika virus infection in human skin fibroblasts. EXCLI J. 2019 Jun 27;18:467–76.

33. Trimarco JD, Heaton BE, Chaparian RR, Burke KN, Binder RA, Gray GC, et al. TMEM41B is a host factor required for the replication of diverse coronaviruses including SARS-CoV-2. PLOS Pathog. 2021 May 27;17(5):e1009599.

34. Eymieux S, Rouillé Y, Terrier O, Seron K, Blanchard E, Rosa-Calatrava M, et al. Ultrastructural modifications induced by SARS-CoV-2 in Vero cells: a kinetic analysis of viral factory formation, viral particle morphogenesis and virion release. Cell Mol Life Sci CMLS. 2021 Jan 15;78(7):3565–76.

35. Cortese M, Lee JY, Cerikan B, Neufeldt CJ, Oorschot VMJ, Köhrer S, et al. Integrative Imaging Reveals SARS-CoV-2-Induced Reshaping of Subcellular Morphologies. Cell Host Microbe. 2020 Dec 9;28(6):853–866.e5.

36. Thompson MR, Sharma S, Atianand M, Jensen SB, Carpenter S, Knipe DM, et al. Interferon γ-inducible Protein (IFI) 16 Transcriptionally Regulates Type I Interferons and Other Interferon-stimulated Genes and Controls the Interferon Response to both DNA and RNA Viruses. J Biol Chem. 2014 Aug 22;289(34):23568–81.

37. Lee S, Lee Y suk, Choi Y, Son A, Park Y, Lee KM, et al. The SARS-CoV-2 RNA interactome. Mol Cell. 2021 Jul 1;81(13):2838–2850.e6.

38. Schmidt N, Lareau CA, Keshishian H, Ganskih S, Schneider C, Hennig T, et al. The SARS-CoV-2 RNA– protein interactome in infected human cells. Nat Microbiol. 2021 Mar;6(3):339–53.

39. Labeau A, Fery-Simonian L, Lefevre-Utile A, Pourcelot M, Bonnet-Madin L, Soumelis V, et al. Characterization and functional interrogation of the SARS-CoV-2 RNA interactome. Cell Rep. 2022 Apr 26;39(4):110744.

40. Murigneux E, Softic L, Aubé C, Grandi C, Judith D, Bruce J, et al. Proteomic analysis of SARS-CoV-2 particles unveils a key role of G3BP proteins in viral assembly. Nat Commun. 2024 Jan 20;15(1):640.

41. Gerke V, Moss SE. Annexins: From Structure to Function. Physiol Rev. 2002 Apr;82(2):331–71.

42. Dreyfuss G, Matunis MJ, Pinol-Roma S, Burd CG. hnRNP PROTEINS AND THE BIOGENESIS OF mRNA. Annu Rev Biochem. 1993 Jul 1;62(Volume 62, 1993):289–321.

43. Luo H, Chen Q, Chen J, Chen K, Shen X, Jiang H. The nucleocapsid protein of SARS coronavirus has a high binding affinity to the human cellular heterogeneous nuclear ribonucleoprotein A1. Febs Lett. 2005 May 9;579(12):2623–8.

44. Shi ST, Huang P, Li H, Lai MMC. Heterogeneous nuclear ribonucleoprotein A1 regulates RNA synthesis of a cytoplasmic virus. EMBO J. 2000 Sep;19(17):4701–11.

45. Wang Y, Zhang X. The Nucleocapsid Protein of Coronavirus Mouse Hepatitis Virus Interacts with the Cellular Heterogeneous Nuclear Ribonucleoprotein A1 *in Vitro* and *in Vivo*. Virology. 1999 Dec 5;265(1):96–109.

46. Beyleveld G, Chin DJ, Olmo EMD, Carter J, Najera I, Cillóniz C, et al. Nucleolar Relocalization of RBM14 by Influenza A Virus NS1 Protein. mSphere [Internet]. 2018 Nov 14 [cited 2026 Jan 2]; Available from: https://journals.asm.org/doi/10.1128/mspheredirect.00549-18

47. Lee N, Yario TA, Gao JS, Steitz JA. EBV noncoding RNA EBER2 interacts with host RNA-binding proteins to regulate viral gene expression. Proc Natl Acad Sci U S A. 2016 Mar 22;113(12):3221–6.

48. Wang X, Tong W, Yang X, Zhai H, Qin W, Liu C, et al. RBM14 inhibits the replication of porcine epidemic diarrhea virus by recruiting p62 to degrade nucleocapsid protein through the activation of autophagy and interferon pathway. J Virol. 2024 Feb 27;98(3):e00182–24.

49. Liu W, Morito D, Takashima S, Mineharu Y, Kobayashi H, Hitomi T, et al. Identification of RNF213 as a Susceptibility Gene for Moyamoya Disease and Its Possible Role in Vascular Development. PLoS ONE. 2011 Jul 20;6(7):e22542.

50. Morito D, Nishikawa K, Hoseki J, Kitamura A, Kotani Y, Kiso K, et al. Moyamoya disease-associated protein mysterin/RNF213 is a novel AAA+ ATPase, which dynamically changes its oligomeric state. Sci Rep. 2014 Mar 24;4:4442.

51. Thery F, Martina L, Asselman C, Zhang Y, Vessely M, Repo H, et al. Ring finger protein 213 assembles into a sensor for ISGylated proteins with antimicrobial activity. Nat Commun. 2021 Oct 1;12(1):5772.

52. Qi Y, Wang M, Jiang Q. PABPC1——mRNA stability, protein translation and tumorigenesis. Front Oncol. 2022 Dec 1;12:1025291.

53. Zhai H, Qin W, Dong S, Yang X, Zhai X, Tong W, et al. PEDV N protein capture protein translation element PABPC1 and eIF4F to promote viral replication. Vet Microbiol. 2023 Sep 1;284:109844.

54. Wang J, Zhang XZ, Sun XY, Tian WJ, Wang XJ. Cellular RNA-binding proteins LARP4 and PABPC1 synergistically facilitate viral translation of coronavirus PEDV. Vet Microbiol. 2024 Nov 1;298:110219.

55. Latifkar A, Levdansky Y, Balabaki A, Nyeo S, Valkov E, Bartel D. mRNA poly(A)-tail length is a battleground for coronavirus–host competition [Internet]. bioRxiv; 2025 [cited 2025 Oct 12]. p. 2025.10.09.680815. Available from: https://www.biorxiv.org/content/10.1101/2025.10.09.680815v1

56. Schattgen SA, Fitzgerald KA. The PYHIN protein family as mediators of host defenses. Immunol Rev. 2011;243(1):109–18.

57. Bosso M, Kirchhoff F. Emerging Role of PYHIN Proteins as Antiviral Restriction Factors. Viruses. 2020 Dec;12(12):1464.

58. Stehlik C. The PYRIN domain in signal transduction. Curr Protein Pept Sci. 2007 Jun;8(3):293–310.

59. Roy A, Ghosh A, Kumar B, Chandran B. IFI16, a nuclear innate immune DNA sensor, mediates epigenetic silencing of herpesvirus genomes by its association with H3K9 methyltransferases SUV39H1 and GLP. Sawyer SL, Akhmanova A, Mahalingam R, editors. eLife. 2019 Nov 4;8:e49500.

60. Howard TR, Lum KK, Kennedy MA, Cristea IM. The Nuclear DNA Sensor IFI16 Indiscriminately Binds to and Diminishes Accessibility of the HSV-1 Genome to Suppress Infection. mSystems. 2022 May 16;7(3):e00198–22.

61. Veeranki S, Choubey D. Interferon-inducible p200-family protein IFI16, an innate immune sensor for cytosolic and nuclear double-stranded DNA: Regulation of subcellular localization. Mol Immunol. 2012 Jan;49(4):567–71.

62. Li T, Diner BA, Chen J, Cristea IM. Acetylation modulates cellular distribution and DNA sensing ability of interferon-inducible protein IFI16. Proc Natl Acad Sci. 2012 Jun 26;109(26):10558–63.

63. Johnstone RW, Kerry JA, Trapani JA. The Human Interferon-inducible Protein, IFI 16, Is a Repressor of Transcription *. J Biol Chem. 1998 Jul 3;273(27):17172–7.

64. Unterholzner L, Keating SE, Baran M, Horan KA, Jensen SB, Sharma S, et al. IFI16 is an innate immune sensor for intracellular DNA. Nat Immunol. 2010 Nov;11(11):997–1004.

65. Jakobsen MR, Bak RO, Andersen A, Berg RK, Jensen SB, Jin T, et al. IFI16 senses DNA forms of the lentiviral replication cycle and controls HIV-1 replication. Proc Natl Acad Sci. 2013 Nov 26;110(48):E4571–80.

66. Jønsson KL, Laustsen A, Krapp C, Skipper KA, Thavachelvam K, Hotter D, et al. IFI16 is required for DNA sensing in human macrophages by promoting production and function of cGAMP. Nat Commun. 2017 Feb 10;8(1):14391.

67. Monroe KM, Yang Z, Johnson JR, Geng X, Doitsh G, Krogan NJ, et al. IFI16 DNA Sensor Is Required for Death of Lymphoid CD4 T Cells Abortively Infected with HIV. Science. 2014 Jan 24;343(6169):428–32.

68. Kaur R, Batra J, Stuchlik O, Reed MS, Pohl J, Sambhara S, et al. Heterogeneous Ribonucleoprotein A1 (hnRNPA1) Interacts with the Nucleoprotein of the Influenza a Virus and Impedes Virus Replication. Viruses. 2022 Feb;14(2):199.

69. Kumar R, Chander Y, Khandelwal N, Nagori H, Verma A, Pal Y, et al. hnRNPA1 regulates early translation to replication switch in SARS-CoV-2 life cycle [Internet]. bioRxiv; 2021 [cited 2025 Oct 12]. p. 2021.07.13.452288. Available from: https://www.biorxiv.org/content/10.1101/2021.07.13.452288v1

70. Schneider WM, Chevillotte MD, Rice CM. Interferon-Stimulated Genes: A Complex Web of Host Defenses. Annu Rev Immunol. 2014;32:513–45.

71. Schoggins JW, Rice CM. Interferon-stimulated genes and their antiviral effector functions. Curr Opin Virol. 2011 Dec 1;1(6):519–25.

72. Tran V, Ledwith MP, Thamamongood T, Higgins CA, Tripathi S, Chang MW, et al. Influenza virus repurposes the antiviral protein IFIT2 to promote translation of viral mRNAs. Nat Microbiol. 2020 Dec;5(12):1490–503.

73. Sullivan OM, Nesbitt DJ, Schaack GA, Feltman EM, Nipper T, Kongsomros S, et al. IFIT3 RNA-binding activity promotes influenza A virus infection and translation efficiency. J Virol. 2025 Jun 11;99(7):e00286–25.

74. Cho KF, Branon TC, Udeshi ND, Myers SA, Carr SA, Ting AY. Proximity labeling in mammalian cells with TurboID and split-TurboID. Nat Protoc. 2020 Dec;15(12):3971–99.

75. Hughes CS, Moggridge S, Müller T, Sorensen PH, Morin GB, Krijgsveld J. Single-pot, solid-phase-enhanced sample preparation for proteomics experiments. Nat Protoc. 2019 Jan;14(1):68–85.

