## Supplementary Figures for "Proximity proteomics reveals a role for IFI16 during human coronavirus infection"

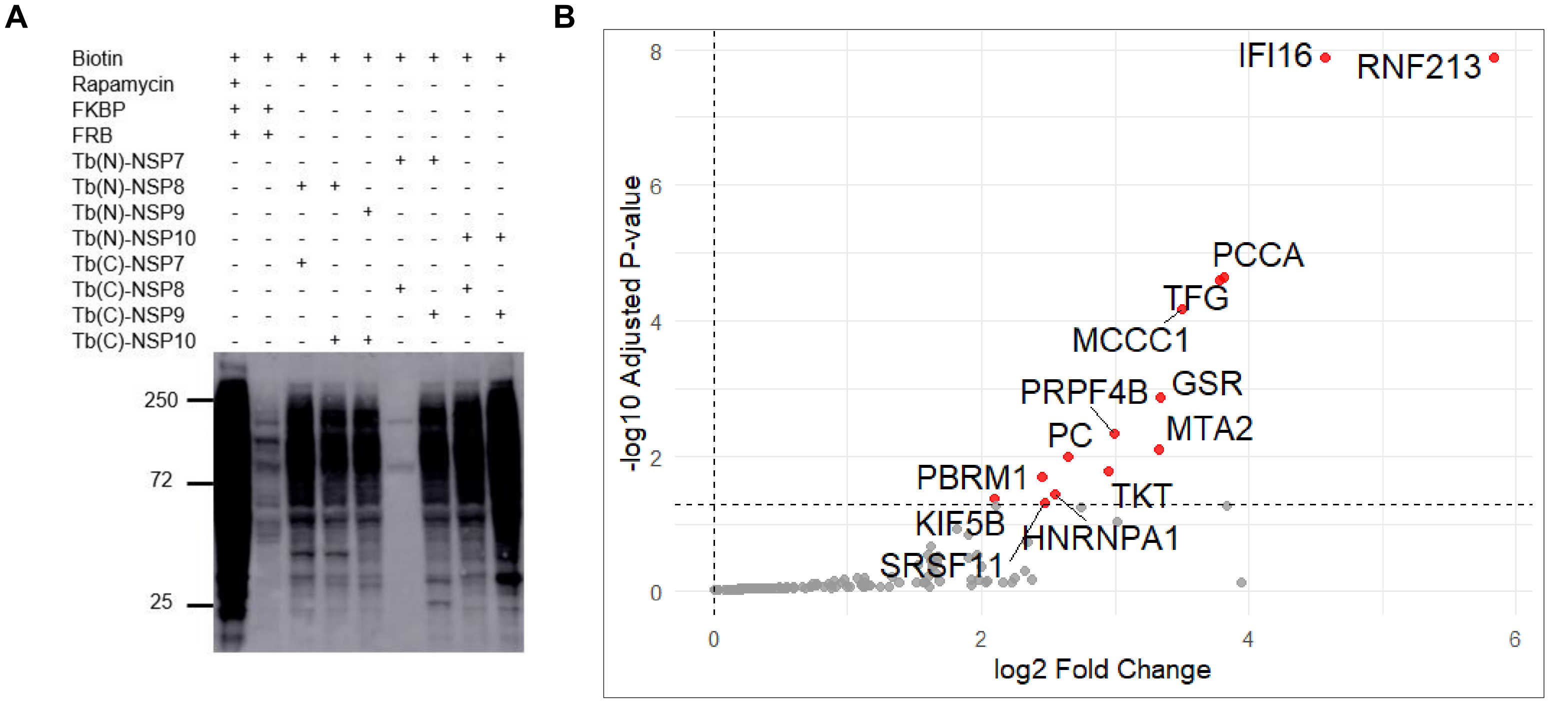


**Supplementary Figure S1. Mass spectrometry identifies proximal partners of SARS-CoV-2 NSP8 and NSP10. A** Western blot to evaluate the formation of functional split-turboID upon transfection of 293T cells with compatible NSP-NSP pairs. **B** Volcano plot showing enriched proteins in clone 2 of A549 cells stably expressing SARS-CoV-2 NSP8-Tb(N) and NSP10- Tb(C).

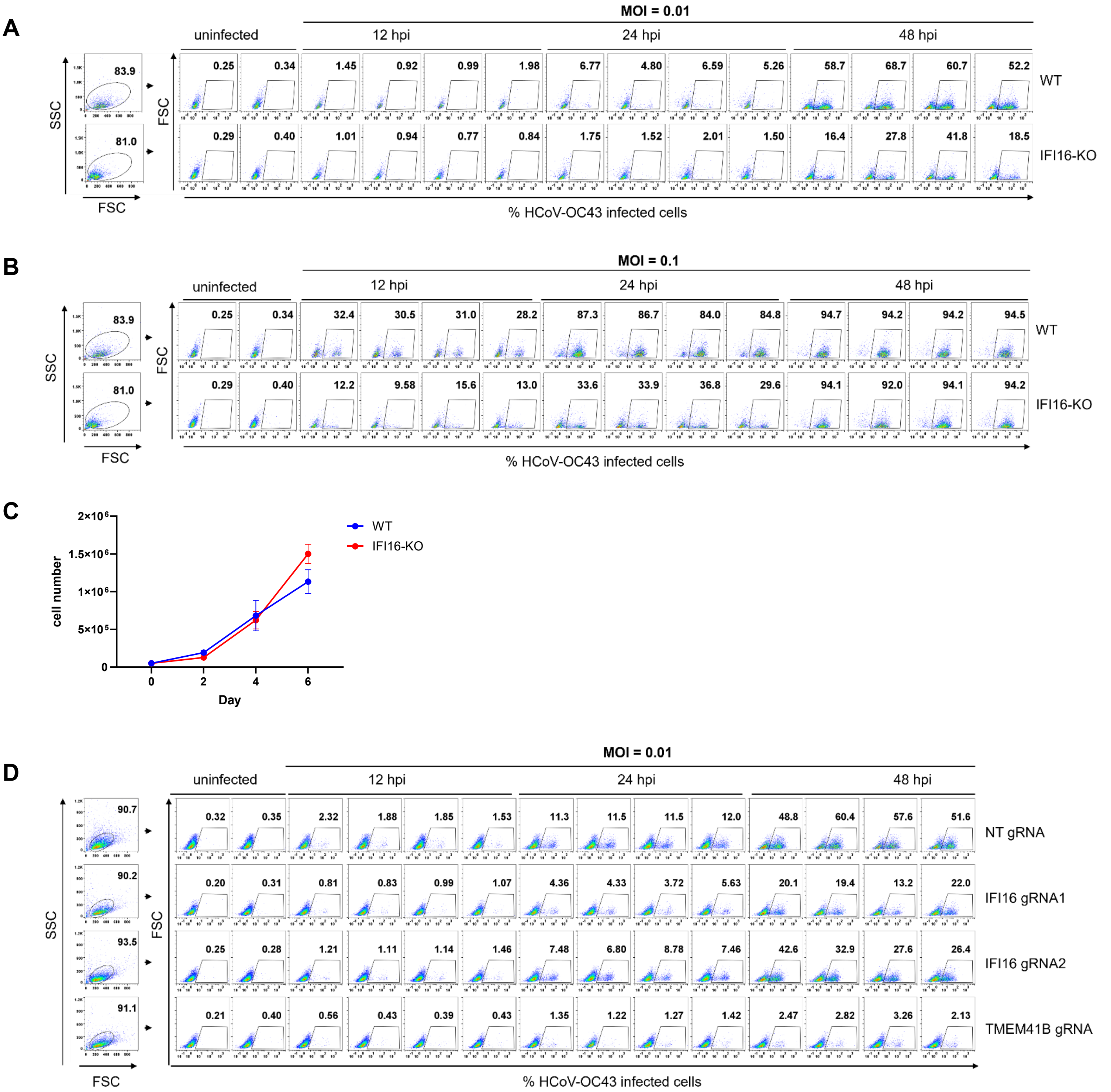


**Supplementary Figure S2. IFI16 promotes coronavirus infection. A** Flow cytometry analysis of WT and IFI16-KO cells infected with HCoV-OC43 at an MOI of 0.01 for 12, 24 and 48 h**. B** Flow cytometry analysis of WT and IFI16-KO cells infected with HCoV-OC43 at an MOI of 0.1 for 12, 24 and 48 h. **C** Cell counts of WT and IFI16-KO cells to determine the impact of IFI16 loss on cell proliferation. Cells were seeded in 12-well plates at 50,000 cells per well (in duplicates) and cell populations were counted with a hemocytometer at the indicated timepoints. **D** Flow cytometry analysis of CRISPRi cell lines infected with HCoV-OC43 at an MOI of 0.01 for 12, 24 and 48 h.

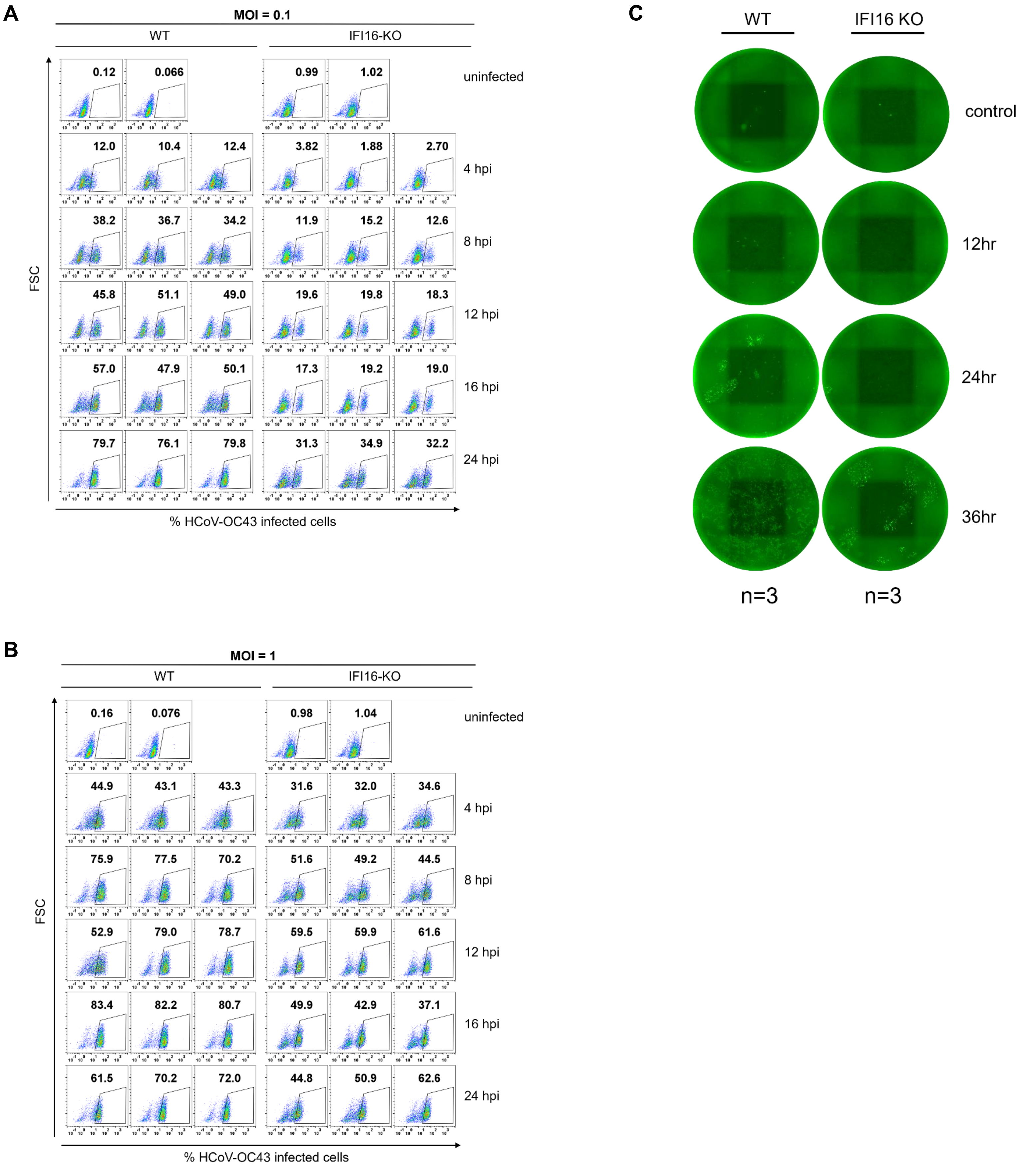
A
MOI = 0.1
WT
IFI16-KO
0.12
0.066
0.99
1.02
uninfected
12.0
10.4
12.4
3.82
1.88
2.70
4 hpi
38.2
36.7
34.2
11.9
15.2
12.6
8 hpi
45.8
51.1
49.0
19.6
19.8
18.3
12 hpi
57.0
47.9
50.1
17.3
19.2
19.0
16 hpi
79.7
76.1
79.8
31.3
34.9
32.2
24 hpi
FSC
% HCoV-OC43 infected cells
B
MOI = 1
WT
IFI16-KO
0.16
0.076
0.98
1.04
uninfected
44.9
43.1
43.3
31.6
32.0
34.6
4 hpi
75.9
77.5
70.2
51.6
49.2
44.5
8 hpi
52.9
79.0
78.7
59.5
59.9
61.6
12 hpi
83.4
82.2
80.7
49.9
42.9
37.1
16 hpi
61.5
70.2
72.0
44.8
50.9
62.6
24 hpi
FSC
% HCoV-OC43 infected cells
C
WT
IFI16 KO
control
12hr
24hr
36hr
n=3
n=3

**Supplementary Figure S3. IFI16 enhances coronavirus replication. A** Time course quantification of infected WT A549 and IFI16-KO cells upon infection with HCoV-OC43 at an MOI = 0.01 (n=3). **B** Time course quantification of infected WT and IFI16-KO cells upon infection with HCoV-OC43 at an MOI = 0.1 (n= 3). **C** Quantification of infectious virus particles produced by WT and IFI16-KO cells upon infection with HCoV-OC43 at an MOI = 0.1 for 12, 24, and 36 h. Supernatant was harvested at each timepoint, and virus titer was determined by titration of harvested supernatant on HCT-8 cells in 96-well plates. Infected cells were stained for HCoV-OC43 N protein and fluorescent foci were imaged using BioTek Cytation C10 (Agilent Technologies).

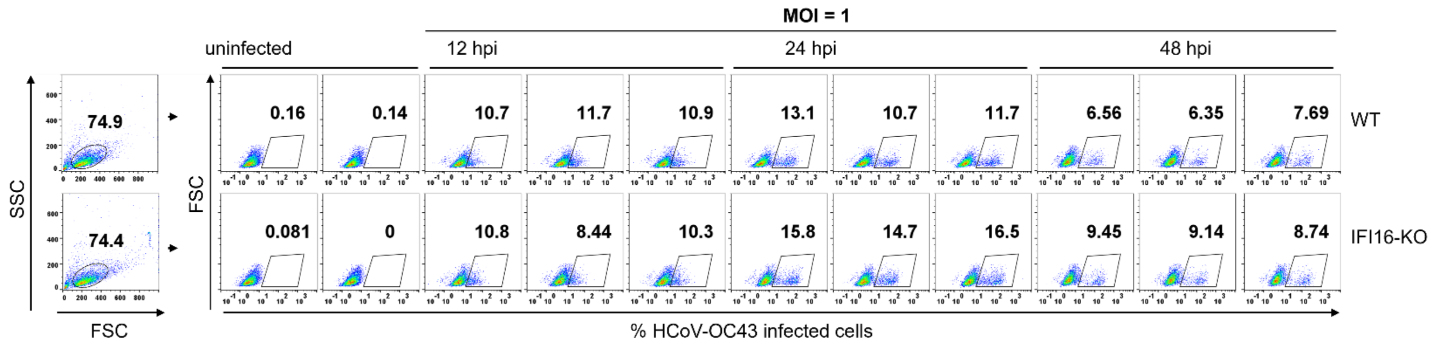


**Supplementary Figure S4. The role of IFI16 in coronavirus replication is independent of its role as a transcriptional regulator of type I IFN.** Flow cytometry was conducted to determine whether the loss of IFI16 abrogates the antiviral response to HCoV-OC43. WT and IFI16-KO cells were pretreated with 200 ng of poly I:C for 24 h, followed by infection with HCoV-OC43 at an MOI =1 for 12, 24 and 48 h.
